# Phased genome sequence of an interspecific hybrid flowering cherry, Somei-Yoshino (*Cerasus × yedoensis*)

**DOI:** 10.1101/573451

**Authors:** Kenta Shirasawa, Tomoya Esumi, Hideki Hirakawa, Hideyuki Tanaka, Akihiro Itai, Andrea Ghelfi, Hideki Nagasaki, Sachiko Isobe

## Abstract

We report the phased genome sequence of an interspecific hybrid, the flowering cherry Somei-Yoshino (*Cerasus* × *yedoensis*). The sequence was determined by single-molecule real-time sequencing technology and assembled using a trio-binning strategy in which allelic variation was resolved to obtain phased sequences. The resultant assembly consisting of two haplotype genomes spanned 690.1 Mb with 4,552 contigs and an N50 length of 1.0 Mb. We predicted 95,076 high-confidence genes, including 94.9% of the core eukaryotic genes. Based on a high-density genetic map, we established a pair of eight pseudomolecule sequences, with highly conserved structures between two genome sequences with 2.4 million sequence variants. A whole genome resequencing analysis of flowering cherry varieties suggested that Somei-Yoshino is derived from a cross between *C. spachiana* and either *C. speciose* or its derivative. Transcriptome data for flowering date revealed comprehensive changes in gene expression in floral bud development toward flowering. These genome and transcriptome data are expected to provide insights into the evolution and cultivation of flowering cherry and the molecular mechanism underlying flowering.

## Introduction

Flowering cherry, called sakura, is Japan’s unofficial national flower and is a popular ornamental tree in Japan and elsewhere. Cherry blossoms are symbols of spring, when blooming typically occurs. Accordingly, flowering cherries are important resources for the tourism industry in the spring season in Japan. More than 200 varieties of flowering cherry are grown (Kato et al. 2012). The nomenclature and, in particular, the genus name (*Prunus* or *Cerasus*) has been under discussion. We use the genus name *Cerasus* in accordance with recent molecular and population genetic analyses (Katsuki and Iketani 2016). Most varieties belong to a species complex with ten basic diploid founders (2n=16), *C. apetala, C. incisa, C. jamasakura, C. kumanoensis, C. leveilleana, C. maximowiczii, C. nipponica, C. sargentii, C. spachiana*, and *C. speciosa*.

Somei-Yoshino (*C.* × *yedoensis*), also known as Yoshino cherry, is the most popular variety of flowering cherry. Somei-Yoshino is believed to have been originally bred in a nursery in the Somei area of Edo (the former name of Tokyo), followed by its spread throughout Japan. Somei-Yoshino is probably derived from an interspecific hybrid between two diploids (2n=16) (Oginuma and Tanaka 1976), *C. spachiana* and *C. speciosa* (Innan et al. 1995; Nakamura et al. 2015a; Takenaka 1963). An alternative hypothesis is that Somei-Yoshino arose from a cross between *C. spachiana* and a hybrid of *C. jamasakura* and *C. speciosa* (Kato et al. 2014). It is self-incompatible, like other members of the Rosaceae family, and accordingly no seeds are produced by self-pollination. Even if self-pollinated seeds are obtained, genotypes would be segregated owing to the high heterozygosity. Therefore, Somei-Yoshino is clonally propagated by grafting or cutting and distributed. The clonality is supported by DNA analyses (Iketani et al. 2007; Innan et al. 1995). Thus, the taxonomic classification has been well investigated. However, to the best of our knowledge, there are few studies of the molecular mechanism underlying flowering in flowering cherry to date, despite extensive analyses of other members of the family Rosaceae.

Some-Yoshino trees are used as standards for forecasting the flowering date of cherry blossoms in the early spring every year. Bud breaking and flowering are important and scientifically intriguing growth stages. In buds, the floral primordia are generally initiated in the summer (late June to August), after which the primordia start to differentiate into floral organs. After differentiation is completed, the buds enter a dormancy period during the winter. Recent studies have evaluated the molecular mechanisms underlying dormancy release as well as flowering in fruit tree species belonging to the family Rosaceae (Lloret et al. 2018; Yamane 2014). Phytohormones and transcriptional regulators involved in dormancy initiation and release have been characterized, including gibberellic acids (GAs) and abscisic acid (ABA). *DELLA* genes, containing a conserved DELLA motif involved in GA signaling, and *CBF/DREB1* (C-repeat-binding factor/dehydration responsive element-binding factor 1) genes involved in cold acclimation have been analyzed in apple (Wisniewski et al. 2015; Yordanov et al. 2014) and Japanese apricot (Lv et al. 2018). The involvement of ethylene signaling, perhaps via crosstalk with ABA, has also been discussed based on a study of *EARLY BUD-BREAK 1* (*EBB1*), which encodes an AP2 type/ethylene-responsive transcription factor (Yordanov et al. 2014). *DORMANCY-ASSOCIATED MADS-BOX* (*DAM*) genes in the same family as *SHORT VEGETATIVE PHASE* (*SVP*) genes (Leida et al. 2010; Yamane et al. 2011), *FLOWERING LOCUS T* (*FT*), and *CENTRORADIALIS* (*CEN*)/*TERMINAL FLOWER 1* (*TFL1*), encoding PEBP-like proteins involved in floral initiation and meristem development, are involved in dormancy (Kurokura et al. 2013). These previous studies provide insight into the genetic basis of dormancy and flowering in fruit tree species belonging to the family Rosaceae.

Genetic and genomic analyses are straightforward approaches to gain insights into the flowering mechanism in cherry blossoms. Whole genome sequences of more than 100 plant species have been published (Michael and VanBuren 2015). Usually, the targets are haploids or inbred lines to simplify the genomic complexity. However, advanced long-read sequencing technologies and bioinformatics methods have made it possible to determine the sequences of complex genomes (Belser et al. 2018; Jiao and Schneeberger 2017; Kyriakidou et al. 2018). For example, an assembly strategy for single-molecule real-time sequencing data has been developed to generate phased sequences in heterozygous regions of F1 hybrids (Chin et al. 2016). Furthermore, chromosome-scale phased genome assemblies for F1 hybrids have been obtained by linked read sequencing technology, providing long-range genome information (Hulse-Kemp et al. 2018), or by single-molecule real-time sequencing combined with Hi-C data (Dudchenko et al. 2017; Kronenberg et al. 2018). Haplotype-resolved sequences have been obtained for F1 cattle by a trio-binning strategy in which genome sequences with allelic variation are resolved before assembly (Koren et al. 2018).

In this study, to determine the molecular mechanisms underlying cherry blossom flowering, we conducted genome and transcriptome analyses of the interspecific hybrid Somei-Yoshino. The genome sequence of another interspecific hybrid flowering cherry, *C.* × *nudiflora*, formerly named *P. yedoensis* (Katsuki and Iketani 2016), has been published (Baek et al. 2018). However, all genomic regions derived from the two different progenitor species (*C. spachiana* and *C. jamasakura*) are totally collapsed. Therefore, we established the phased genome sequence of *C.* × *yedoensis*, Somei-Yoshino, representing the two genomes of the probable progenitors (*C. spachiana* and *C. speciosa*). Using the genome sequences as a reference, a time-course transcriptome analysis of Somei-Yoshino floral buds and flowers, with a special focus on dormancy and flowering-related genes, was also conducted to characterize the physiological changes during flowering.

## Materials and methods

### Plant materials

A Somei-Yoshino tree grown in Ueno Park (Tokyo, Japan) was used for genome assembly. This tree, i.e., #136, is presumed to be the original according to a polymorphism analysis of three genes and its location (Nakamura et al. 2015a; Nakamura et al. 2015b). In addition, 139 varieties, including a Somei-Yoshino clone maintained at Shimane University (SU), Shimane, Japan, were used for a genetic diversity analysis (Supplementary Table S1). An F1 mapping population, YSF1, was produced by hand pollination between Yama-Zakura and another clone of Somei-Yoshino as a female and male parent, respectively, both of which are planted at the Kazusa DNA Research Institute (KDRI), Chiba, Japan. The Somei-Yoshino clones at SU and KDRI were used for the transcriptome analysis.

### Clustering analysis of genetically divergent varieties

Genomic DNAs of the 139 varieties were extracted from young leaves using the DNeasy Plant Mini Kit (Qiagen, Hilden, Germany) and double-digested with the restriction enzymes *Pst*I and *Eco*RI. ddRAD-Seq libraries were constructed as described previously (Shirasawa et al. 2016) and sequenced using the Illumina HiSeq2000 (San Diego, CA, USA) to obtain 93 bp paired-end reads. Low-quality reads were trimmed using PRINSEQ v. 0.20.4 (Schmieder and Edwards 2011) and adapter sequences were removed using fastx_clipper (parameter, -a AGATCGGAAGAGC) in FASTX-Toolkit v. 0.0.13 (http://hannonlab.cshl.edu/fastx_toolkit). The high-quality reads were mapped onto genome sequences of either *P. avium* (Shirasawa et al. 2017), *P. mume* (Zhang et al. 2012), or *P. persica* (International Peach Genome et al. 2013) using Bowtie2 v. 2.2.3 (Langmead and Salzberg 2012). Biallelic SNPs were called from the mapping results using the mpileup command in SAMtools v. 0.1.19 (Li et al. 2009), and low-quality SNPs were removed using VCFtools v. 0.1.12b (Danecek et al. 2011) with the following criteria: including only sites with a minor allele frequency of ≥0.05 (--maf 0.05), including only genotypes supported by ≥5 reads (--minDP 5), including only sites with a quality value of ≥999 (--minQ 999), and excluding sites with ≥50% missing data (--max-missing 0.5). A dendrogram based on the SNPs was constructed using the neighbor-joining method implemented in TASSEL 5 (Bradbury et al. 2007) and population structure was investigated using ADMIXTURE v. 1.3.0 with default settings (*K* = 1 to 20) (Alexander et al. 2009).

### Assembly of the ‘Somei-Yoshino’ genome

Genomic DNA was extracted from young leaves of Somei-Yoshino tree #136 using the DNeasy Plant Mini Kit (Qiagen). A paired-end sequencing library (insert size of 500 bp) and three mate-pair libraries (insert sizes of 2 kb, 5 kb, and 8 kb) were constructed using the TruSeq PCR-free Kit (Illumina) and Mate-pair Kit (Illumina), respectively, and sequenced using the MiSeq and HiSeqX platforms (Illumina). The size of the Somei-Yoshino genome was estimated using Jellyfish v. 2.1.4 (Marcais and Kingsford 2011). High-quality reads after removing adapter sequences and trimming low-quality reads as described above were assembled using SOAPdenovo2 v. 1.10 (Luo et al. 2012) (parameter -K 121). Gaps, represented by Ns in the sequence, were filled with high-quality paired-end reads using GapCloser v. 1.10 (Luo et al. 2012) (parameter -p 31). The resultant sequences were designated CYE_r1.0.

High-molecular-weight DNA was extracted from young leaves of ‘Somei-Yoshino’ tree #136 using Genomic Tip (Qiagen) to prepare the SMRTbell library (PacBio, Menlo Park, CA, USA). The sequence reads obtained from the PacBio Sequel system were assembled using FALCON-Unzip (Chin et al. 2016) to obtain an assembly, CYE_r2.0. Furthermore, the PacBio reads were divided into two subsets using the TrioCanu module of Canu v. 1.7 (Koren et al. 2018), in which Illumina short reads of two probable ancestors of Somei-Yoshino, i.e., *C. spachiana* ‘Yaebeni-shidare’ and *C. speciosa* ‘Ohshima-zakura,’ were employed. Each subset of data was assembled and polished using FALCON assembler v. 2.1.2 (Chin et al. 2013). The two assemblies were designated CYEspachiana_r3.0 and CYEspeciosa_r3.0, and were combined to obtain CYE_r3.0, representing the Somei-Yoshino genome. Assembly completeness was evaluated using BUSCO v. 3.0.2 (Simao et al. 2015), for which Plants Set was employed as datasets, and a mapping rate analysis of whole genome sequence data for Somei-Yoshino reads to the references was performed (see below for details).

### Genetic map construction and pseudomolecule establishment

Genomic DNA was extracted from the ovules of YSF1 seeds using the Favorgen Plant Kit (Ping-Tung, Taiwan) and digested with *Pst*I and *Eco*RI to construct the ddRAD-Seq library. The library was sequenced on the Illumina NextSeq platform. High-quality reads were mapped onto CYEspaciana_r3.0 and CYEspeciosa_r3.0 using Bowtie2 v. 2.2.3 (Langmead and Salzberg 2012). Biallelic SNPs were called from the mapping results using the mpileup command in SAMtools v. 0.1.19 (Li et al. 2009), and low-quality SNPs were deleted using VCFtools v. 0.1.12b (Danecek et al. 2011) with the criteria used for the clustering analysis described above. The SNPs from the two references were merged, grouped, and ordered using Lep-Map3 v. 0.2 (Rastas 2017). Flanking sequences of the SNP sites (100 bases up-and downstream of the SNPs) were compared with the genome sequence of sweet cherry, PAV_r1.0 (Shirasawa et al. 2017), by BlastN with a cutoff value of 1E-40. Probable misassemblies found in the mapping process were broken, and the resultant sequence set was designated CYE_r3.1. According to map positions, the CYE_r3.1 sequences were oriented and assigned to the genetic map of ‘Somei-Yoshino’ to establish pseudomolecule sequences. Sequence variation between the two pseudomolecule sequences, CYEspaciana_r3.1 and CYEspeciosa_r3.1, was detected using the show-snps function of MUMMER v. 3.23 (Kurtz et al. 2004), for which outputs from NUCmer were employed. In parallel, the genome structure of CYE_r3.1_pseudomolecule was compared with those of sweet cherry, peach, Japanese apricot, and apple using D-Genies (Cabanettes and Klopp 2018).

### Gene prediction and annotation

Total RNA was extracted from 12 stages of buds within 1 month in 2017 as well as from leaves, stems, sepals, petals, stamens, and carpels. RNA-Seq libraries were prepared using the TruSeq Stranded mRNA Sample Preparation Kit (Illumina) and sequenced by MiSeq. The obtained reads were mapped to the CYE_r3.1 sequences to determine gene positions using TopHat2 v. 2.0.14 (Kim et al. 2013). The positional information was used in BREAKER2 v. 2.1.0 (Hoff et al. 2016) to gain training data sets for AUGUSTUS v. 3.3 (Stanke et al. 2006) and GeneMark v. 4.33 (Lomsadze et al. 2005). The two training sets and a preset of SNAP v. 2006-07-28 for *Arabidopsis* as well as peptide sequences of *P. avium* (v1.0.a1), *P. persica* (v2.0.a1), and *Malus* × *domestica* (GDDH13 v1.1) registered in the Genome Database for Rosaceae (Jung et al. 2019) and those of *P. mume* (Zhang et al. 2012) were analyzed using MAKER pipeline v. 2.31.10 (Cantarel et al. 2008) to predict putative protein-coding genes in the CYE_r3.1 sequences. Genes annotated using Hayai-Annotation Plants v. 1.0 (Ghelfi et al. 2019) (with a sequence identity threshold of 80% and query coverage of 80%) were selected as a high-confidence gene set.

### Gene clustering, multiple sequence alignment, and divergence time estimation

Potential orthologues were identified from genes predicted in seven genomes (two genomes of Somei-Yoshino and one each of *P. avium, P. mume, P. persica*, and *M.* × *domestica*, as well as *Arabidopsis thaliana* as an outgroup) using OrthoMCL v. 2.0.9 (Li et al. 2003). The single copy orthologues in the seven genomes were used to generate a multiple sequence alignment using MUSCLE v. 3.8.31 (Edgar 2004), in which indels were eliminated by Gblocks v. 0.91b (Castresana 2000). A phylogenetic tree based on the maximum-likelihood algorithm was constructed from the alignments with the Jones-Taylor-Thornton model in MEGA X v. 10.0.5 (Kumar et al. 2018). The divergence time was calculated using MEGA X v. 10.0.5 (Kumar et al. 2018) assuming that the divergence time between *M.* × *domestica* and *P. persica* was approximately 34 to 67 MYA in TIMETREE (Kumar et al. 2017).

### Repetitive sequence analysis

A database of repeat sequences of the Somei-Yoshino genome was established using RepeatModeler v. 1.0.11 (Smit et al. 2008-2015). The repeat database as well as that registered in Repbase (Bao et al. 2015) were used to predict repetitive sequences in CYE_r3.1 using RepeatMasker v. 4.0.7 (Smit et al. 2013-2015).

### Whole genome resequencing analysis

Genomic DNA of eight representative lines of the SU collection and one of the parental lines of the mapping population, Yama-Zakura, were digested with NEBNext dsDNA Fragmentase (New England BioLabs, Ipswich, MA, USA) for whole genome shotgun library preparation using the Illumina TruSeq PCR-free Kit. The sequences were determined on the Illumina NextSeq platform. Read trimming, read mapping to the CYE_r3.1 sequence, and SNP identification were performed as described above. Effects of SNPs on gene functions were evaluated using SnpEff v. 4.2 (Cingolani et al. 2012).

### Transcriptome analysis

Additional RNA-Seq libraries were prepared from buds at 24 stages collected in 2017 at KDRI and in 2014 and 2015 at SU using the TruSeq Stranded mRNA Library Prep Kit (Illumina) and sequenced on the NextSeq500 (Illumina). High-quality reads after removing adapter sequences and trimming low-quality reads as mentioned above were mapped to the pseudomolecule sequences of CYE_r3.1 using HISAT2 v. 2.1.0 (Kim et al. 2015), and reads on each gene model were quantified and normalized to determine FPKM values using StringTie v. 1.3.5 (Pertea et al. 2015) and Ballgown v.2.14.1 (Frazee et al. 2015) in accordance with the protocol paper (Pertea et al. 2016). The R package WGCNA v.1.66 (Langfelder and Horvath 2008) was used for network construction and module detection.

## Results

### Clustering analysis of cherry varieties

We obtained approximately 1.9 million (M) high-quality reads per line after trimming adapters and low-quality sequences from the ddRAD-Seq library. The reads were mapped onto the genome sequences of *P. avium* (PAV_r1.0), *P. mume*, and *P. persica* (v1.0) with mapping alignment rates of 70.8%, 77.8%, and 68.7%, respectively (Supplementary Table S2). We detected 46,278 (*P. avium*), 31,973 (*P. mume*), and 33,199 (*P. persica*) high-confidence SNPs. A clustering tree based on the 46,278 SNPs and a population structure analysis indicated that the cherry collection was derived from at least eight founders (*K* = 8) (Supplementary Figure S1). The Somei-Yoshino genome consisted of *C. spachiana* and *C. speciose* genomic features.

### Assembly of the Somei-Yoshino genome

The ‘Somei-Yoshino’ genome size was estimated by a k-mer analysis with 14.3 Gb of paired-end reads (20.7×) obtained by MiSeq (Supplementary Table S3). The distribution of distinct k-mers (k = 17) showed two peaks at multiplicities of 18 and 37, indicating heterozygous and homozygous regions, respectively (Supplementary Figure S2). This result suggested that the heterozygosity of the Somei-Yoshino genome was high. In other words, Somei-Yoshino is likely an interspecific hybrid harboring components of two different genomes. The total size of the two genomes was approximately 690 Mb.

Totals of 132.5 Gb of paired-end reads (192× genome coverage) and 69.1 Gb of mate-pair data (100×) (Supplementary Table S3) were assembled into 1.2 million scaffold sequences. The total length of the resultant scaffolds, i.e., CYE_r1.0, was 686.9 Mb, including 63.6 Mb of Ns with an N50 length of 142.5 kb (Supplementary Table S4). Only 62.3% of complete single copy orthologs in plant genomes were identified in a BUSCO analysis (Supplementary Table S4). Paired-end reads of Somei-Yoshino (20.7×) were mapped onto CYE_r1.0 with a mapping rate of 76.6%. We found that 82.4% of SNPs were homozygous for the reference type. Ideally, both rates should be close to 100% if the assembly was fully extended and the two genomes were separated, or phased. Distributions of the sequence depth of coverage showed a single peak at the expected value of 21× (Supplementary Figure S3). When we mapped the reads to the sequence of *C.* × *nudiflora* (Pyn.v1) (Baek et al. 2018), two peaks at 22× (expected) and 44× (double the expected value) were observed (Supplementary Figure S3), indicating a mixture of phased and unphased sequences.

To extend the sequence contiguity and to improve the genome coverage, PacBio long-read technology was employed to obtain 37.3 Gb of reads (54×) with an N50 read length of 17 kb (Supplementary Table S3). The long reads were assembled using FALCON-Unzip into 3,226 contigs [470 primary contigs (488 Mb) and 2,756 haplotigs (116 Mb)] spanning a total length of 605.4 Mb with an N50 length of 2.3 Mb, i.e., CYE_r2.0 (Supplementary Table S4). A BUSCO analysis indicated that 97.0% of complete BUSCOs (9.1% single copy and 87.9% duplicated, as expected) were represented in the assembly (Supplementary Table S4). The mapping rate of the Somei-Yoshino reads was 95.3%, and 97.1% of SNPs were homozygous for the reference type. Most of the sequences were phased, with one major peak of genome coverage at 21× (Supplementary Figure S3); however, the total length was 13% shorter than the estimated size and no haplotype information was available.

We used a trio-binning approach to obtain the entire sequences of the two haplotype sequences. The long reads (37.3 Gb, 54×) were divided into two subsets based on whole genome resequencing of the two lines, i.e., *C. spachiana* (Yaebini-shidare) and *C. speciose* (Ohshima-zakura). The resultant subsets included 18.9 Gb and 18.2 Gb for *C. spachiana* and *C. speciosa*, respectively, and 0.3 Mb of unassigned reads. The subsets were separately assembled to obtain 2,281 contigs (717 primary contigs and 1,564 associated contigs including duplicated repetitive sequences) covering 350.1 Mb, i.e., CYEspachiana_r3.0, and 2,271 contigs (800 primary contigs and 1,471 associated contigs) covering 340.0 Mb, i.e., CYEspachiana_r3.0 (Supplementary Table S4). The total sequence (i.e., CYE_r3.0) spanned 690.1 Mb and consisted of 4,552 contigs with an N50 length of 1.0 Mb (Supplementary Table S4). The complete BUSCO score for CYE_r3.0 was 96.8% (10.6% single copy and 86.2% duplicated, as expected), while those for CYEspachiana_r3.0 and CYEspeciosa_r3.0 were 90.9% (69.3% single copy and 21.6% duplicated) and 88.9% (72.1% single copy and 16.8% duplicated), respectively (Supplementary Table S4). The mapping rate of the Somei-Yoshino reads was as high as 96.3%, and 96.2% of SNPs were homozygous for the reference type. The sequence depth of coverage was distributed as expected, with a single peak at 20× (Supplementary Figure S3). Therefore, CYE_r3.0 was used for further analyses because it satisfied all of the established criteria.

### Genetic map for Somei-Yoshino

Approximately 2.0 million high-quality ddRAD-Seq reads per sample were obtained from YSF1 and mapped to either CYEspachiana_r3.0 or CYEspeciosa_r3.0 with alignment rates of 79.3% and 80.3%, respectively (Supplementary Table S5). We detected 16,145 and 17,462 SNPs from the alignments with the references of CYEspachiana_r3.0 and CYEspeciosa_r3.0, respectively. Of these, 23,532 heterozygous SNPs in ‘Somei-Yoshino’ were used for a linkage analysis. The SNPs were assigned to eight groups and ordered, covering 458.8 cM with 16,933 SNPs in 694 genetic bins (Supplementary Tables S6 and S7). The map was split into two for *C. spachiana* and *C. speciosa*, covering 448.9 cM with 8,280 SNPs (628 genetic bins) and 446.3 cM with 8,653 SNPs (645 genetic bins), respectively. The genetic bins were common for 579 loci on the two maps, suggesting that the sequences in the common bins were the same loci. A comparison of the genetic maps with the genome sequence of sweet cherry, PAV_r1.0 (Supplementary Figure S4), indicated a high similarity of the genome structures in the two species.

### Genetic anchoring of the assemblies to the chromosomes

In the genetic mapping process, we found 19 potential misassemblies in 18 contig sequences of CYE_r3.0. The contigs were broken between SNPs mapped to different linkage groups. Finally, we obtained 4,571 contigs with an N50 length of 918.2 kb and the same total length (690.1 Mb). This final version of contigs was named CYE_r3.1, consisting of CYEspachiana_r3.1 (2,292 contigs, N50 length of 1.2 Mb) and CYEspeciosa_r3.1 (2,279 contigs, N50 length of 800.6 kb) (Table 1). Of these, 184 CYEspachiana_r3.1 contigs (221.8 Mb) and 262 CYEspeciosa_r3.1 contigs (199.2 Mb) were assigned to the genetic maps (Supplementary Tables S8). The contigs were connected with 10,000 Ns to establish the Somei-Yoshino pseudomolecule sequences consisting of 4,571 contigs covering 418 Mb. The structures of the two pseudomolecule sequences were well conserved (Fig. 1). We observed 2,371,773 and 2,392,937 sequence variants, including SNPs and indels, in CYEspachiana_r3.1 (one variant every 93 bp) and CYEspeciosa_r3.1 (one variant every 83 bp), respectively, of which 0.4% were deleterious mutations (Supplementary Tables S9). The structure of the Somei-Yoshino genome showed high synteny with the genomes of other members of Rosaceae (Supplementary Figure S5).

**Table 1.**
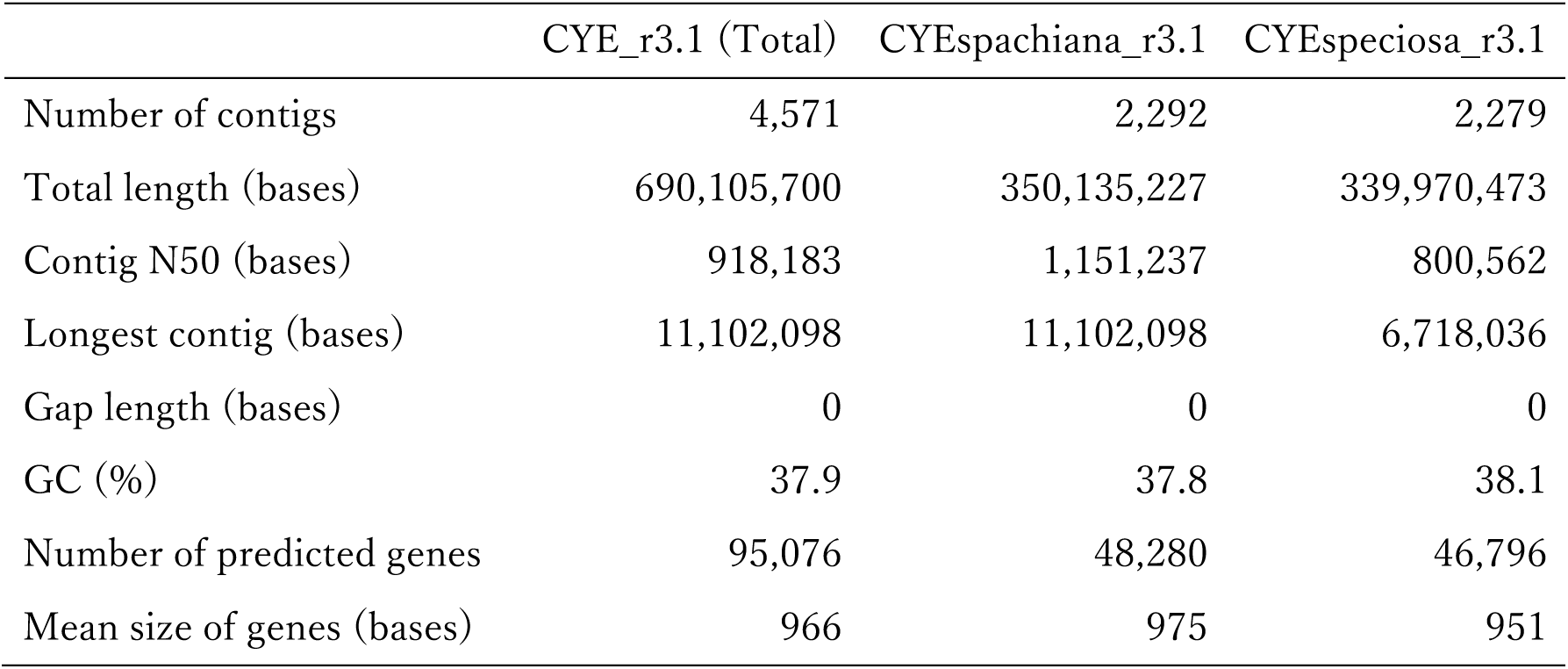
Assembly statistics of the final version of the Somei-Yoshino genome sequence

**Figure 1.**
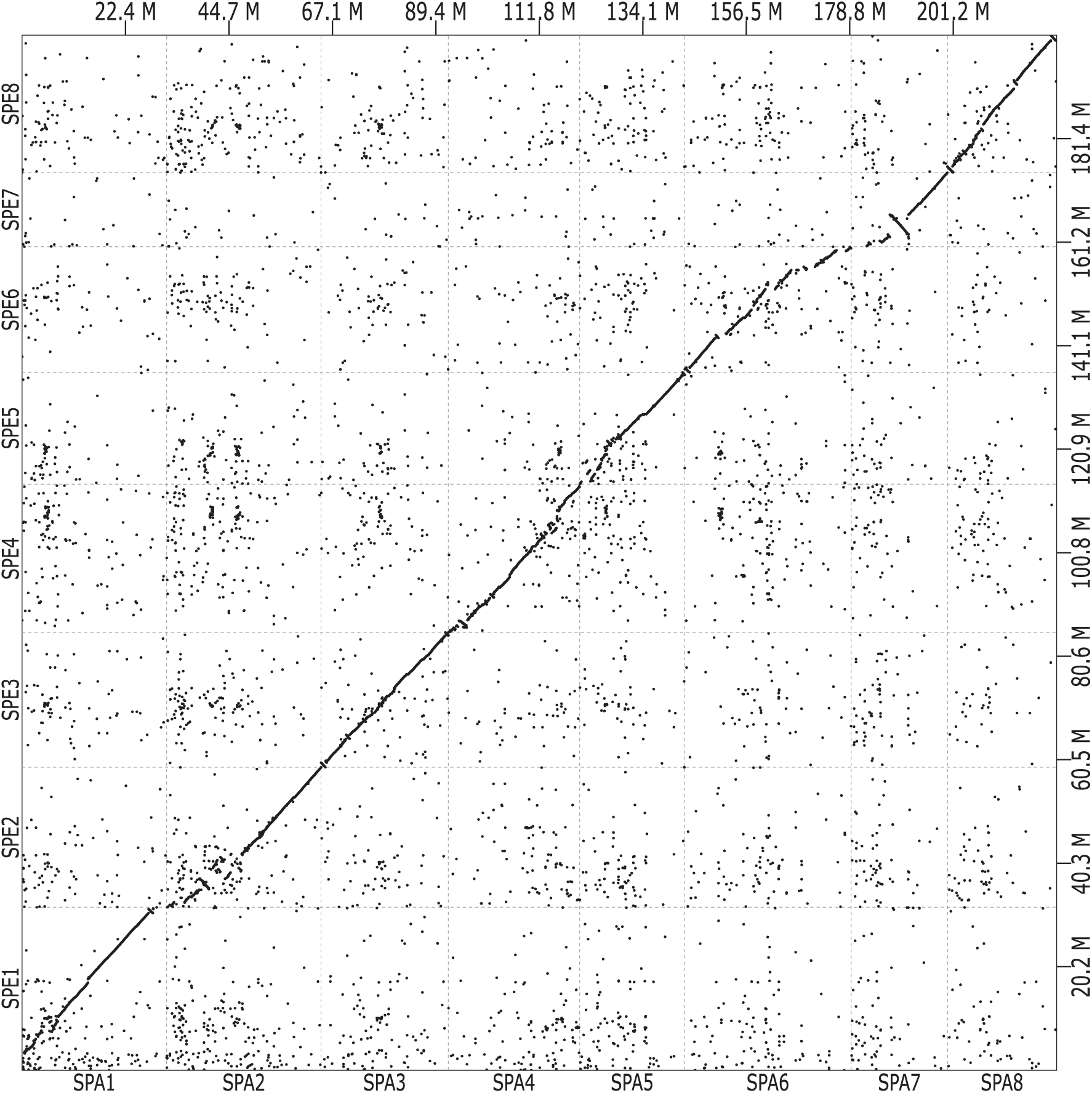
Synteny of the two haplotype pseudomolecule sequences of the Somei-Yoshino genome. *X*-and *Y*-axis are sequences of CYE_r3.1spachiana (SPA1 to 8) and CYE_r3.1speciosa (SPE1 to 8), respectively.

### Gene prediction and annotation

We initially predicted 222,168 putative genes using the MAKER pipeline. All genes were annotated by a similarity search against the UniProtKB database using the Hayai-Annotation Plants pipeline to select 94,776 non-redundant high-confidence genes. Then, 300 genes showing sequence similarity to genes involved in flowering and dormancy in the family Rosaceae (Supplementary Table S10) were manually added. A total of 95,076 genes (48,280 and 46,796 from CYEspachiana_r3.1 and CYEspeciosa_r3.1, respectively) were selected as a high-confidence gene set for CYE_r3.1 (Table 1). The total length of the coding sequences was 91.9 Mb (13.3% of the CYE_r3.1) with an N50 length of 1,512 bases and a GC content of 44.8%. This gene set included 94.9% complete BUSCOs (12.8% single copy and 82.1% duplicated). Out of the 95,076 genes, 26,463 (27.8%), 34,996 (36.8%), and 46,502 (48.9%) were assigned to Gene Ontology slim terms in the biological process, cellular component, and molecular function categories, respectively (Supplementary Table S11). Furthermore, 3,972 genes had enzyme commission numbers.

We found two pairs of self-incompatible genes, *S* determinants for pollen (S-RNase) and pistils (SFB: *S* haplotype-specific F-box); CYE_r3.1SPE0_g058440.1 (S-RNase) and CYE_r3.1SPE0_g058430.1 (SFB) were *S* genes of the *PyS1* haplotype, and CYE_r3.1SPE0_g046700.1 (S-RNase) and CYE_r3.1SPE0_g046660.1 (SFB) were *S* genes of *PyS2*. For dormancy, we detected a cluster of six *DAM*-like genes, as reported in the Japanese apricot genome (Zhang et al. 2012), in the pseudomolecule sequence of SPA1 (CYE_r3.1SPA1_g039840.1 to CYE_r3.1SPA1_g039890.1). In addition, *CBF* gene clusters were also found in SPA5 (CYE_r3.1SPA5_g014520.1 to CYE_r3.1SPA5_g014610.1) and SPE5 (CYE_r3.1SPE5_g016380.1 to CYE_r3.1SPE5_g016430.1).

### Divergence time of Somei-Yoshino ancestors

The predicted genes were clustered with those of apple, sweet cherry, Japanese apricot, peach, and *Arabidopsis* to obtain 29,091 clusters, involving 36,396 and 35,559 genes from CYEspachiana_r3.1 and CYEspeciosa_r3.1, respectively (Supplementary Table S12). Among these, 8,125 clusters were common across the tested species, and 1,254 consisting of one gene from each genome were selected for divergence time estimation. When the divergence time between apple and peach was set to 34 to 67 MYA, the divergence time between the two haplotype sequences of Somei-Yoshino was set to 5.52 MYA (Figure 2).

**Figure 2.**
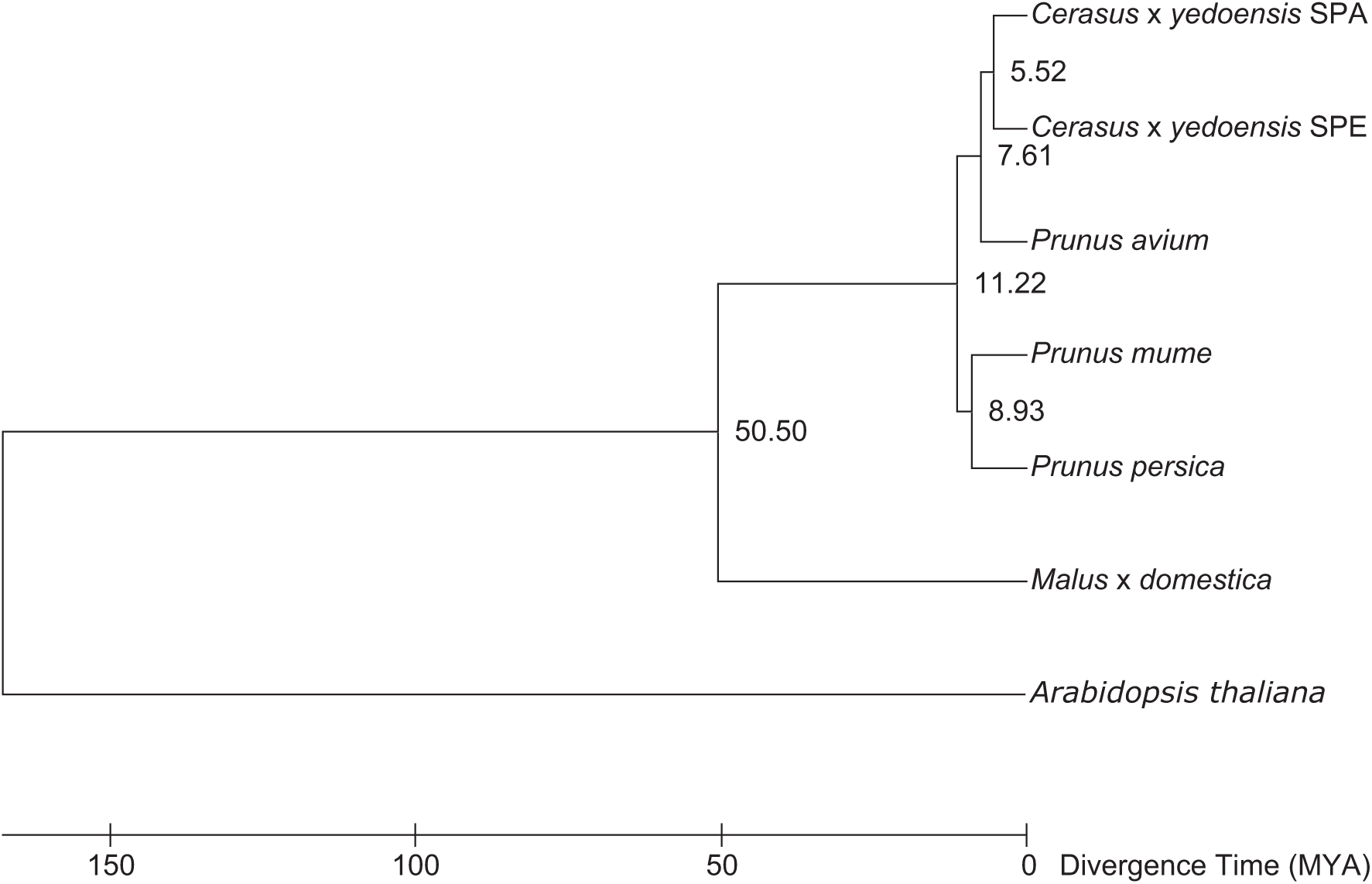
Phylogenetic tree indicating the divergence time of Somei-Yoshino. The two genomes of Somei-Yoshino are indicated by SPA and SPE, representing *C. spachiana* and *C. speciosa*, respectively. Divergence times (MYA; million years ago) between branches are shown.

### Repetitive sequence analysis

A total of 293.3 Mb (42.5%) of CYE_r3.1 (690.1 Mb) was identified as repetitive sequences, including transposable elements (Supplementary Table S13), which occupied 142.9 Mb (40.8%) and 150.4 Mb (44.2%) of CYEspachiana_r3.1 and CYEspeciosa_r3.1, respectively. The most prominent repeat types were long-terminal repeat retrotransposons (104.0 Mb; 14.1%), e.g., *Gypsy* -and *Copia*-types, followed by DNA transposons (65.1 Mb; 8.8%).

### Whole genome resequencing analysis

Approximately 136 million high-quality whole genome sequence reads was obtained from eight representatives in a population structure analysis (Supplementary Table S14) and the parents of the mapping population, Yama-Zakura and ‘Somei-Yoshino.’ In addition, 250 million sequence reads of *C.* × *nudiflora* (Baek et al. 2018) (SRA accession number SRX3900230) was also employed. The reads were aligned to CYE_r3.1 as a reference with a mapping rate of 88.0%, on average. From the alignment data, we detected 2,307,670 SNPs and 169,664 indels, including 658,873 SNPs and 42,286 indels (28.3%) in CYEspachiana_r3.1 and 1,648,797 SNPs and 127,378 indels (71.7%) in CYEspeciosa_r3.1. Of these, 8,872 SNPs (0.4%) were deleterious mutations (Supplementary Tables S15).

In Somei-Yoshino, the reads were evenly mapped to the references of CYEspachiana_r3.1 (48.7%) and CYEspeciosa_r3.1 (47.6%) (Supplementary Figure S6). Most of the loci (94.5% of SNPs in CYEspachiana_r3.1 and 96.9% in CYEspeciosa_r3.1) were homozygous for the reference type, as expected (Supplementary Figure S7). Only 61.7% and 52.9% of SNPs in *C.* × *nudiflora* were reference-type homozygotes on CYEspachiana_r3.1 and CYEspeciosa_r3.1, respectively (Supplementary Figure S6), and read mapping rates were 52.2% (CYEspachiana_r3.1) and 39.8% (CYEspeciosa_r3.1) (Supplementary Figure S7).

In *C. spachiana* (Yaebeni-shidare), 69.8% of the reads were preferentially mapped to CYEspachiana_r3.1 (Supplementary Figure S6), and 80.1% of SNPs detected in CYEspachiana_r3.1 were homozygous for the reference type (Supplementary Figure S7). In *C. speciose* (Ohshima-zakura), 61.1% of reads were mapped to CYEspeciosa_r3.1 (Supplementary Figure S6) and 73.5% of SNPs in CYEspeciosa_r3.1 were homozygous for the reference type (Supplementary Figure S7). In the remaining seven lines, mapping rates on CYEspeciosa_r3.1 were higher than those on CYEspachiana_r3.1, as in *C. speciose* (Ohshima-zakura) (Supplementary Figure S6).

### Transcriptome analysis of flowering dates

RNA-Seq reads were obtained from 12 stages of buds collected every month from May 2014 to April 2015 (Supplementary Table S16) as well as from the 12 stages from 2 to 34 days before anthesis in 2017 used for gene prediction. After trimming, the reads as well as those for the six organs used in the gene prediction analyses were mapped to CYE_r3.1 with a mapping rate of 67.6%, on average. Among the 95,076 predicted genes, 72,248 (76.0%) with a variance across samples of ≥1 were selected. A WGCNA analysis was performed with the expression data for the 24 buds to generate 31 highly co-expressed gene clusters, referred to as modules (Supplementary Figure S8). The modules were roughly grouped into three main classes expressed in the previous year of flowering, within 1 month, and within 1 week (Supplementary Figure S9).

Based on the literature and databases for Rosaceae, we identified dormancy-and flowering-associated genes (i.e., *DELLA, CBF*/*DREB1, EBB1, DAM* (*SVP*), *FT*, and *CEN*/*TFL1* genes). We detected 35 predicted genes in the Somei-Yoshino genome, 16 of which were expressed in ≥1 sample. The expression patterns basically agreed with those of the modules and could be roughly classified into five groups (Figure 3). The first group (blue and magenta gene modules in Supplementary Figure S8) consisted of four genes homologous to *DELLA* genes. Their expression levels were elevated in the floral buds about 1 month before anthesis; expression was also observed in young vegetative buds. The second group (turquoise, brown, and salmon gene modules) were highly expressed in the summer and autumn (from July to November) in the floral buds. Six genes homologous to *CBF*/*DREB1* belonged to this group; however, these were classified into three different clusters on the dendrogram. The third group (turquoise gene module) consisted of two *EBB1* homologs and one *DAM* (*SVP*) homolog; these genes were highly expressed in the autumn and winter (from October to December). In the fourth group (turquoise gene module), genes were highly expressed in the winter 2–3 months before anthesis and were homologous to *FT* genes. The fifth group (red gene module) solely included *CEN*/*TFL1*-like genes specifically expressed in vegetative state buds before flower differentiation.

**Figure 3.**
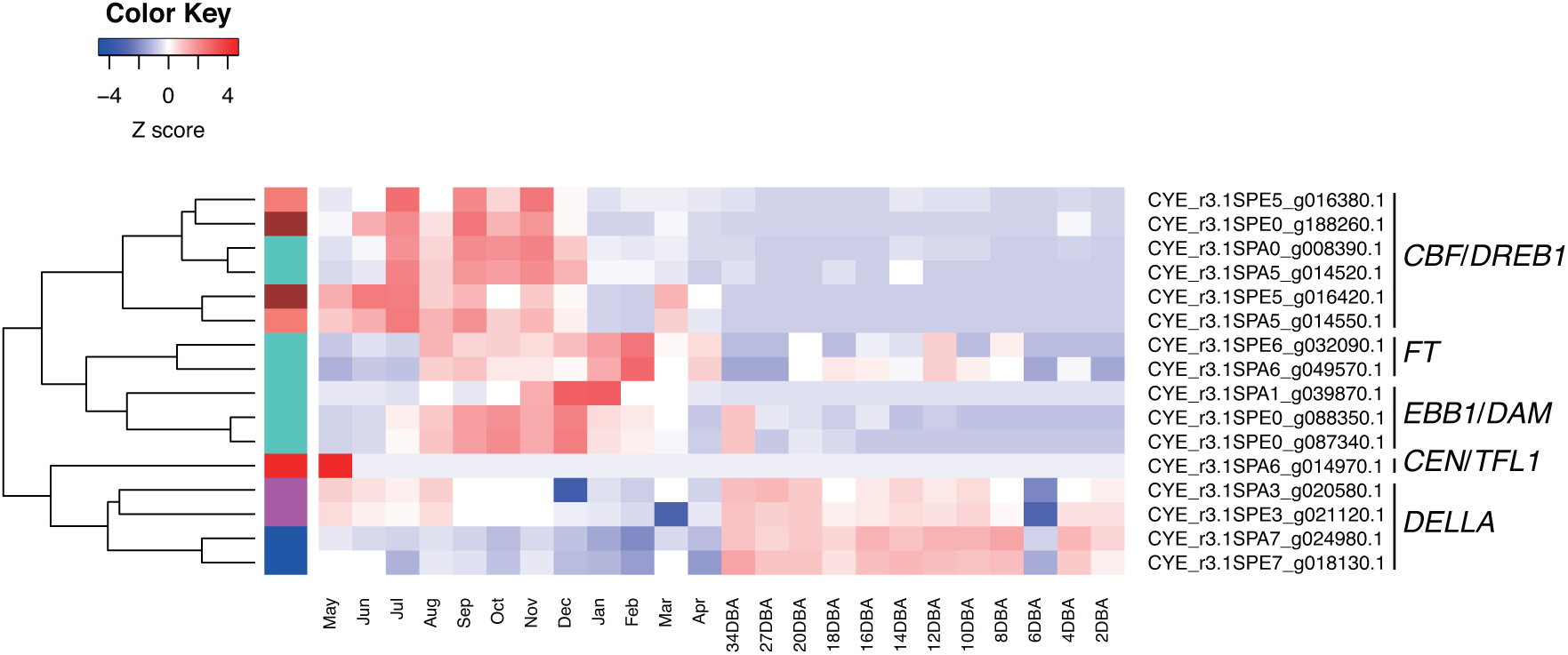
Heat map representing expression patterns of dormancy and flowering genes in Somei-Yoshino buds. Colors in each block represent a continuum of gene expression levels with Z-score-transformed FPKM (low-to-high gene expression levels are represented by blue to red). May to Apr are the months and 34DBA to 2DBA are days before anthesis when bud samples were collected. Gene modules based on WGCNA (see also Supplementary Figure S8) are shown as colored bars between the dendrogram and heatmap.

## Discussion

We obtained the genome sequence of the flowering cherry Somei-Yoshino. To the best of our knowledge, this is the first report of a phased genome sequence of an interspecific hybrid in the family Rosaceae or in the kingdom of Plantae, broadly, although genome sequences have been reported for several species belonging to Rosaceae (Jung et al. 2019). Although the genome of another interspecific hybrid cherry flower, *C.* × *nudiflora*, has been reported (Baek et al. 2018), the two homoeologous ancestral genomes (*C. spachiana* and *C. jamasakura*) are totally collapsed, as indicted by the double peaks of sequence depth (Supplementary Figure S3), resulting in a short assembly size (323.8 Mb). The genome complexity of interspecific hybrids could be compared to those of polyploids and highly heterozygous species. Genome sequences of polyploids and F1 hybrids have been obtained (Chin et al. 2016; Hulse-Kemp et al. 2018) by single-molecule real-time sequencing technology, linked read sequencing, optical maps, and Hi-C (Belser et al. 2018; Jiao and Schneeberger 2017; Kyriakidou et al. 2018). These technologies to obtain phased genome assemblies are limited by haplotype switching (Kronenberg et al. 2018), where two haplotypes are patched to make mosaic genome sequences.

We employed the trio-binning technique (Koren et al. 2018) to determine haplotype phases before assembly. This technique was initially developed to construct phased genome sequences of an F1 hybrid between cattle subspecies. Since sequence reads of two sub-genomes were divided into two subsets according to the sequences of the parents, haplotype switching is avoidable. We applied the trio-binning technique to the interspecific hybrid cherry tree. We verified the quality and accuracy of the resultant assembly, CYE_r3.0, by a BUSCO analysis (Supplementary Table S4), the mapping rate of Somei-Yoshino reads to the assemblies (Supplementary Figure S6), and SNP genotypes detected in the mapping results. In addition, the genetic map (Supplementary Table S6 and S7) and a comparative analysis of the pseudomolecule sequences (Figure 1 and Supplementary Figure S4 and S5) also supported the quality and accuracy of the assembly. The results of this study suggested that the trio-binning strategy is useful for determining phased genome sequences for highly heterozygous genomes of interspecific hybrids.

Our genome data provided insight into the progenitors of Somei-Yoshino. Our results were consistent with the conclusions of Baek et al. (2018), who found that Somei-Yoshino, *C.* × *yedoensis*, is distinct from a variety in Jeju Island, Korea, *C.* × *nudiflora*. In the present study, a population structure analysis indicated that Somei-Yoshino was established by two founders, *C. spachiana* and *C. speciosa* (Figure 2, Supplementary Figure S1), as suggested in previous studies (Innan et al. 1995; Takenaka 1963). In a whole genome resequencing analysis, sequence reads of *C. spachiana* ‘Yaebeni-shidare’ were preferentially mapped to SPA sequences (Supplementary Figure S6), and genotypes of most SNPs were homozygous for the reference type (Supplementary Figure S7). This indicated that the sequence similarity of *C. spachiana* ‘Yaebeni-shidare’ and CYEspachiana_r3.1 was high and therefore that *C. spachiana* is a candidate parent. While reads of *C. speciosa* ‘Ohshima-zakura’ were mapped to CYEspeciosa_r3.1 sequences (Supplementary Figure S6), the frequency of SNP genotypes homozygous for the reference type was not as high as that for *C. spachiana* (Supplementary Figure S7). This observation suggests that *C. speciosa* is not an actual parent of Somei-Yoshino (Kato et al. 2014). Somei-Yoshino genome data can be used in future studies of the origin to determine the most likely parents.

We obtained a number of predicted genes. Transcriptome data for the developing bud provided a comprehensive overview of genes expressed during dormancy and flowering processes (Figure 3). Our analysis was based on previous studies of key genes and fundamental molecular mechanisms underlying dormancy (Lloret et al. 2018; Yamane 2014). Despite some discrepancies, the gene expression patterns observed in our study were generally consistent with previously observed patterns in deciduous fruit tree species in Rosaceae. The relatively high expression levels of *DELLA* genes observed at 1 month before anthesis corresponded to the time at which the bud typically transitions from endodormancy to ecodormancy (Lv et al. 2018). GA signaling may reactivate bud development internally at the ecodormancy stage (Wen et al. 2016). The relatively high expression levels of *CBF*/*DREB1* in the summer and decreased expression levels toward the winter is consistent with a role in cold acclimation, as previously reported in almond (Saibo et al. 2012). We detected one *DAM* gene that was highly expressed in dormant buds in the winter, in agreement with previous reports (Yamane et al. 2006); however, two *EBB1* genes, assigned to the same module as *DAM* genes, showed different expression patterns from those in apple and poplar, in which the genes exhibit sharp increases in expression before bud breaking (Wisniewski et al. 2015; Yordanov et al. 2014). This inconsistency may be explained by differences in regulatory mechanisms underlying bud breaking. *FT* genes showed elevated expression levels in buds in February, when endodormancy is almost completed. In addition to the function of floral induction, unknown functions of *FT* genes during dormancy are possible. Interestingly, transgenic plum (*Prunus domestica*) with overexpressed poplar *FT* (*PtFT1*) does not enter a state of endodormancy upon cold treatment or, alternatively, has no chilling requirement after dormancy is established (Srinivasan et al. 2012). Further studies of the role of *FT* genes in dormancy are needed. *CEN*/*TFL1* was highly expressed only in vegetative buds before floral initiation. This observation was consistent with other previous results for species in the family Rosaceae (Esumi et al. 2010; Mimida et al. 2009). Our transcriptome data for flowering cherry successfully revealed the comprehensive changes in gene expression during floral bud development toward flowering. The expression patterns of above genes in this study and supposed regulation network for dormancy release of woody plants (Falavigna et al. 2019; Lloret et al. 2018; Singh et al. 2018) are jointly summarized in Figure 4. The transcriptome data set provides a basis for further research aimed at identifying additional genes involved in floral bud development and flowering. Especially, identifying genes involved in the regulation of flowering under *FT* gene (protein) signaling and GA signaling processes is intrigued, and those may be able to utilize for accurate forecasting the flowering date of cherry blossoms.

**Figure 4.**
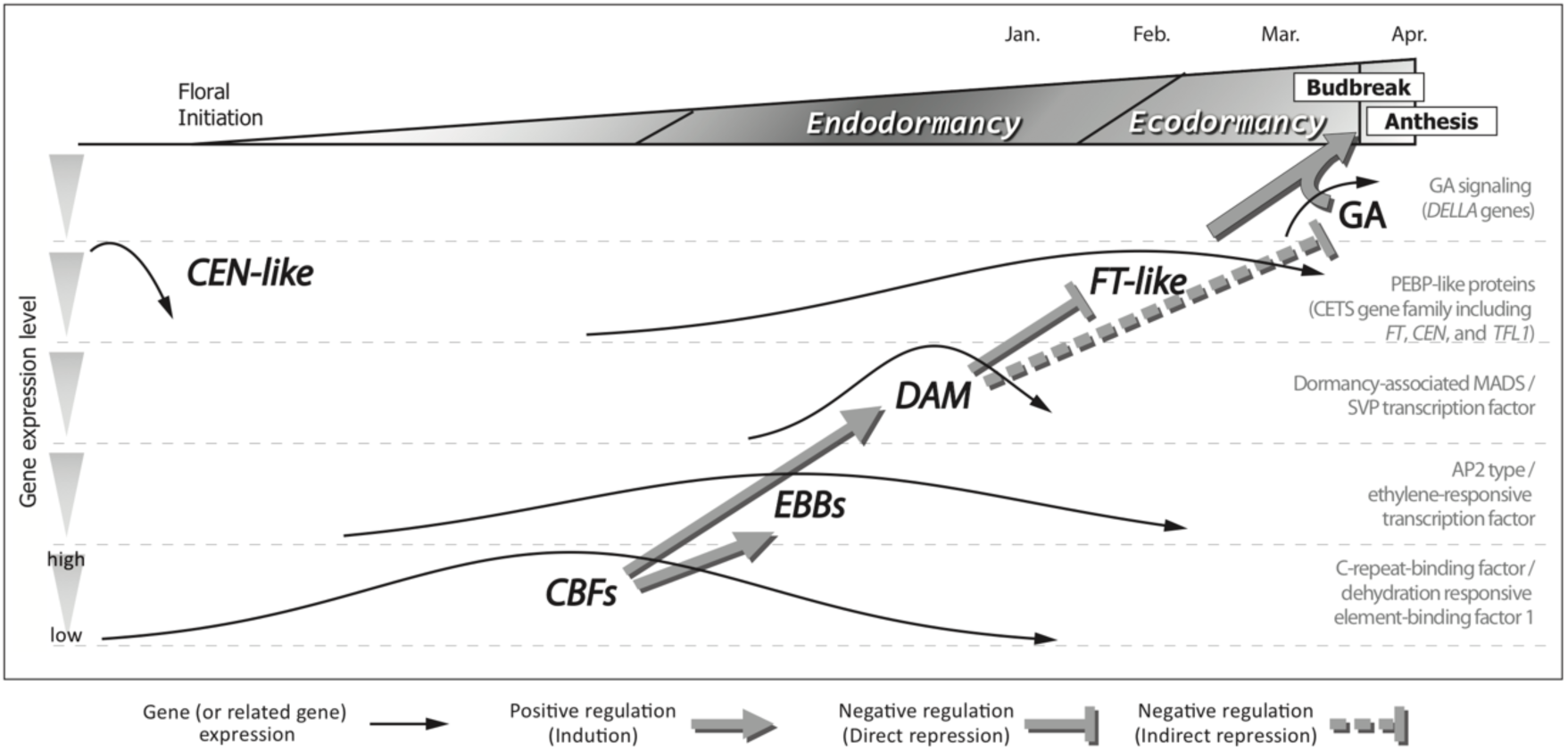
A putative regulation model for dormancy release and flowering with expression patterns of related genes in Somei-Yoshino buds. The supposed regulation mechanism for dormancy and flowering is based on recent studies and reviews in woody plants (Falavigna et al. 2019; Lloret et al. 2018; Singh et al. 2018). The gene expression patterns represented as black arrows are based on Figure 3.

The genome and transcriptome data obtained in this study are expected to accelerate genomic and genetic analyses of flowering cherry. Owing to the complicated genomes, it is necessary to build additional *de novo* assemblies for divergent flowering cherries, which is a challenging task. Genome-graph-based pan-genome analyses could be used to characterize the complex genomes (Rakocevic et al. 2019). The Somei-Yoshino genome sequence would be a resource for the flowering cherry pan-genome analyses. It may provide insights into the evolution and cultivation of flowering cherry as well as the molecular mechanism underlying flowering traits in the species and in the family Rosaceae, and it may guide the future cultivation and breeding of flowering cherry.

## Data availability

The sequence reads are available from the DDBJ Sequence Read Archive (DRA) under the accession numbers DRA008094, DRA008096, DRA008097, DRA008099, and DRA008100. The WGS accession numbers of assembled scaffold sequences are BJCG01000001-BJCG01004571 (4,571 entries). The genome assembly data, annotations, gene models, genetic maps, and DNA polymorphism information are available at DBcherry (http://cherry.kazusa.or.jp).

## Acknowledgments

We thank Ueno Park (Tokyo, Japan) for providing the Somei-Yoshino sample. We are grateful to Drs G. Concepcion and P. Peluso (PacBio, CA, USA) and Mr. K. Osaki (Tomy Digital Biology, Tokyo, Japan) for their helpful advice, and S. Sasamoto, S. Nakayama, A. Watanabe, T. Fujishiro, Y. Kishida, C. Minami, A. Obara, H. Tsuruoka, and M. Yamada (Kazusa DNA Research Institute) for their technical assistance. This work was supported by the Kazusa DNA Research Institute Foundation, and supported in part by a Grant-in-Aid for Young Scientists (B) No. 26850017 (to T. E.) from Japan Society for the Promotion of Science (JSPS).

