## Supplementary Figure for "Phased genome sequence of an interspecific hybrid flowering cherry, Somei-Yoshino (*Cerasus × yedoensis*)"

**Supplementary Table S1** Genetically divergent lines of flowering cherry used in this study
**Supplementary Table S2** Sequence reads for the ddRAD-Seq analysis of flowering cherry lines
**Supplementary Table S3** Number of whole genome shotgun reads for Somei-Yoshino
**Supplementary Table S4** Assembly statistics for the Somei-Yoshino genome
**Supplementary Table S5** Sequence reads in the ddRAD-Seq analysis for the F1 mapping population
**Supplementary Table S6** Genetic map of Somei-Yoshino
**Supplementary Table S7** SNP loci on the genetic map
**Supplementary Table S8** Summary of the Somei-Yoshino pseudomolecule sequence
**Supplementary Table S9** Number of sequence variants between the two genomes of Somei-Yoshino and impacts on gene functions
**Supplementary Table S10** Genes underlying flowering and dormancy in Rosaceae
**Supplementary Table S11** Gene IDs, annotations, and expression levels for predicted genes
**Supplementary Table S12** Gene clusters based on genes in six species in the family Rosaceae and *Arabidopsis thaliana*
**Supplementary Table S13** Repetitive sequences in the Somei-Yoshino genome
**Supplementary Table S14** Number of whole genome shotgun reads of genetically divergent lines of flowering cherry
**Supplementary Table S15** Number of sequence variants among 11 genetically divergent lines
**Supplementary Table S16** Number of RNA-Seq reads for Somei-Yoshino

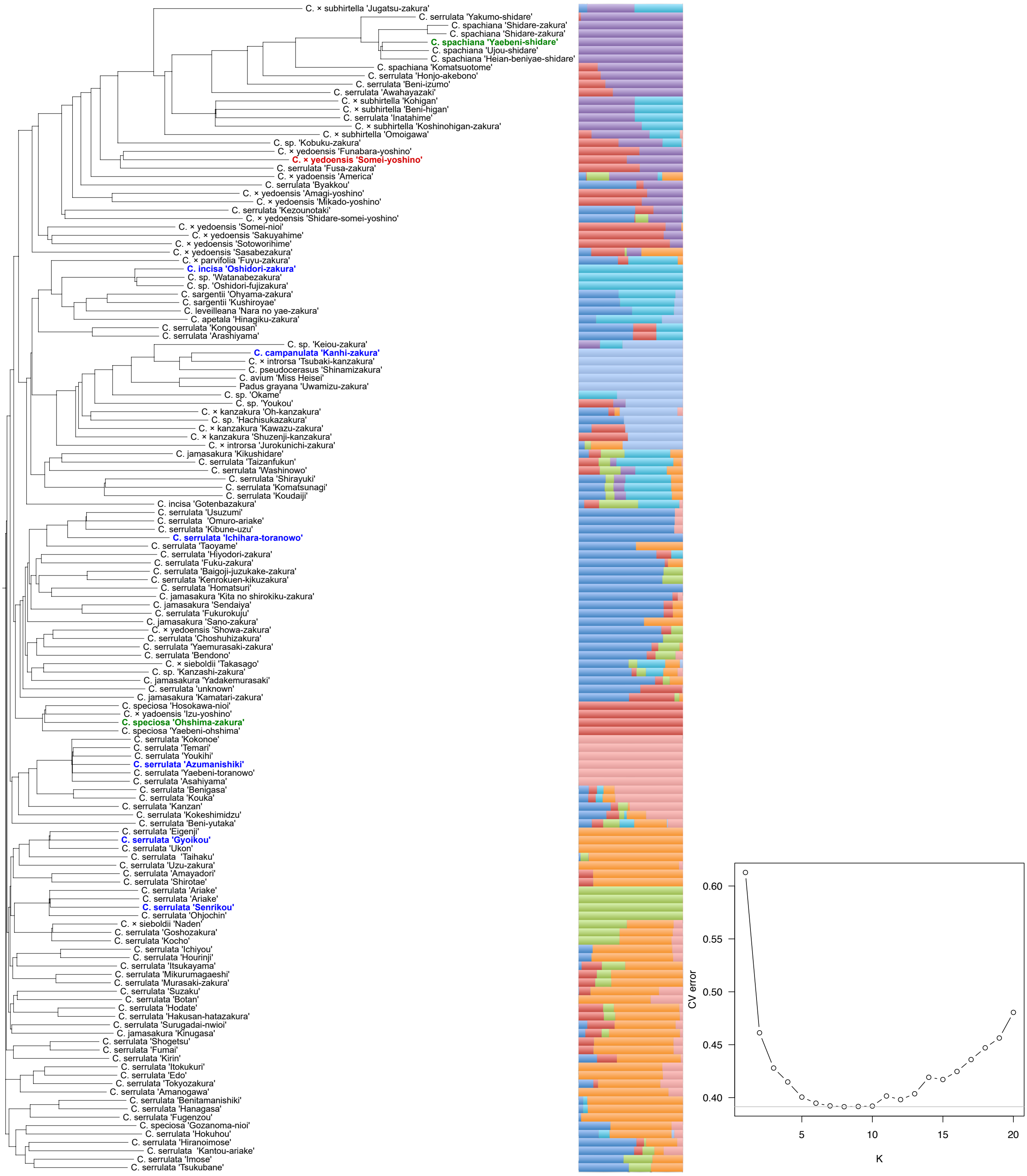


**Supplementary Figure S1.** Dendrogram and population structure analysis of flowering cherries

The dendrogram is based on genetic distances calculated by the neighbor-joining method. Somei-Yoshino is shown in red, and varieties selected for whole genome resequencing are in blue, including two possible ancestral lines shown in green. Line chart indicates cross-validation errors in the admixture analysis. The best fit model is K = 8.

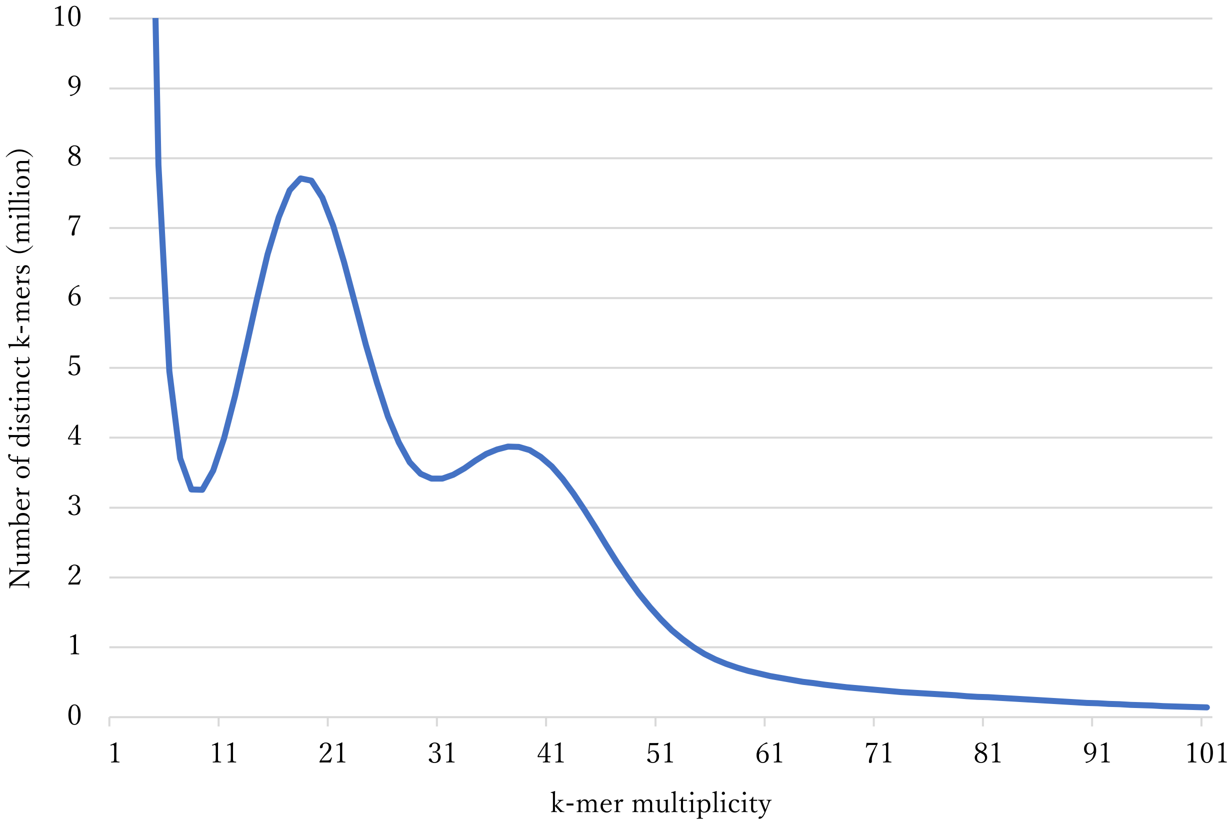


**Supplementary Figure S2.** Genome size estimation for Somei-Yoshino with the distribution of the number of distinct k-mer (k = 17) with the given multiplicity values

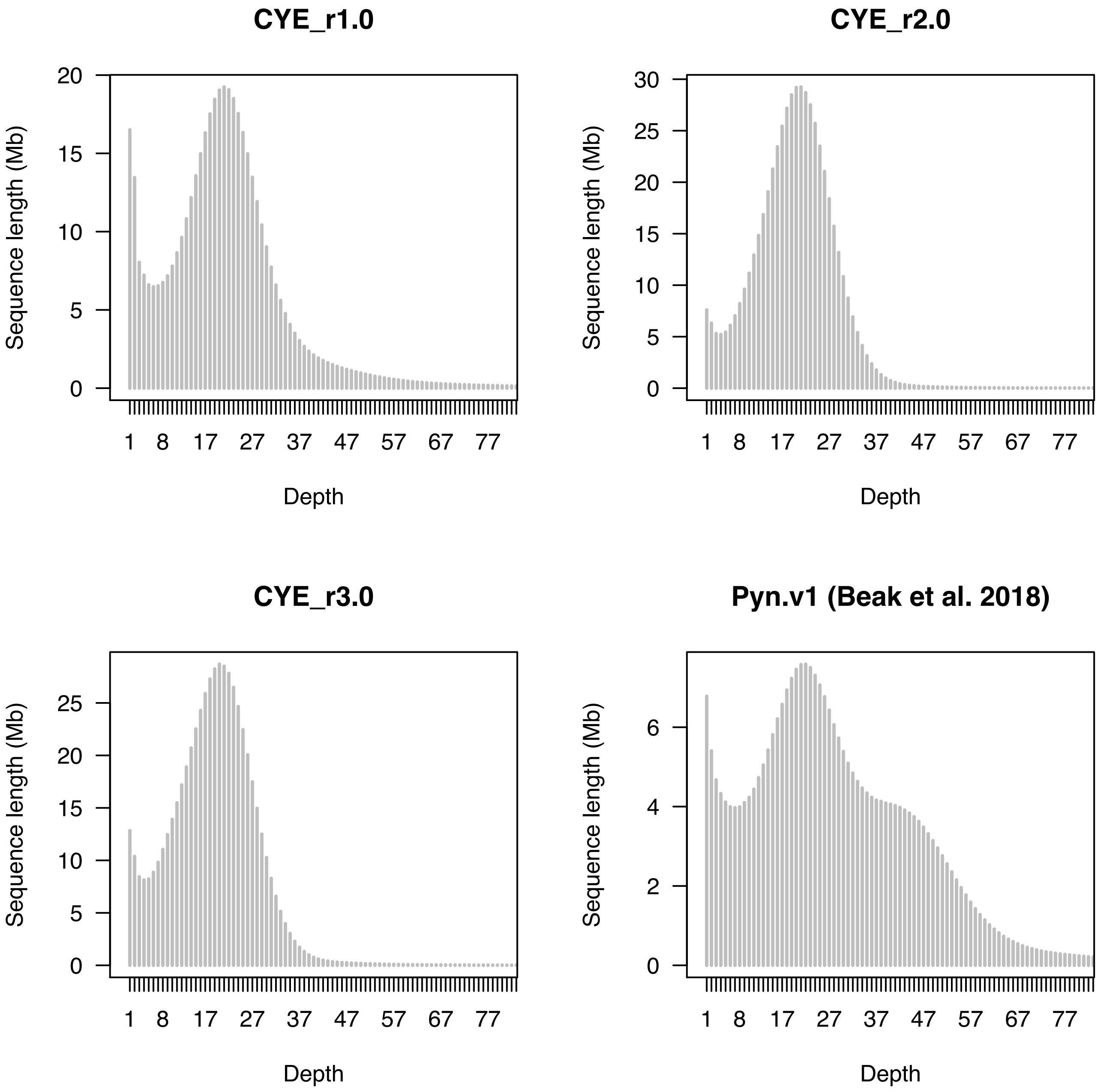


**Supplementary Figure S3.** Distribution of depths of Somei-Yoshino reads on genome assemblies

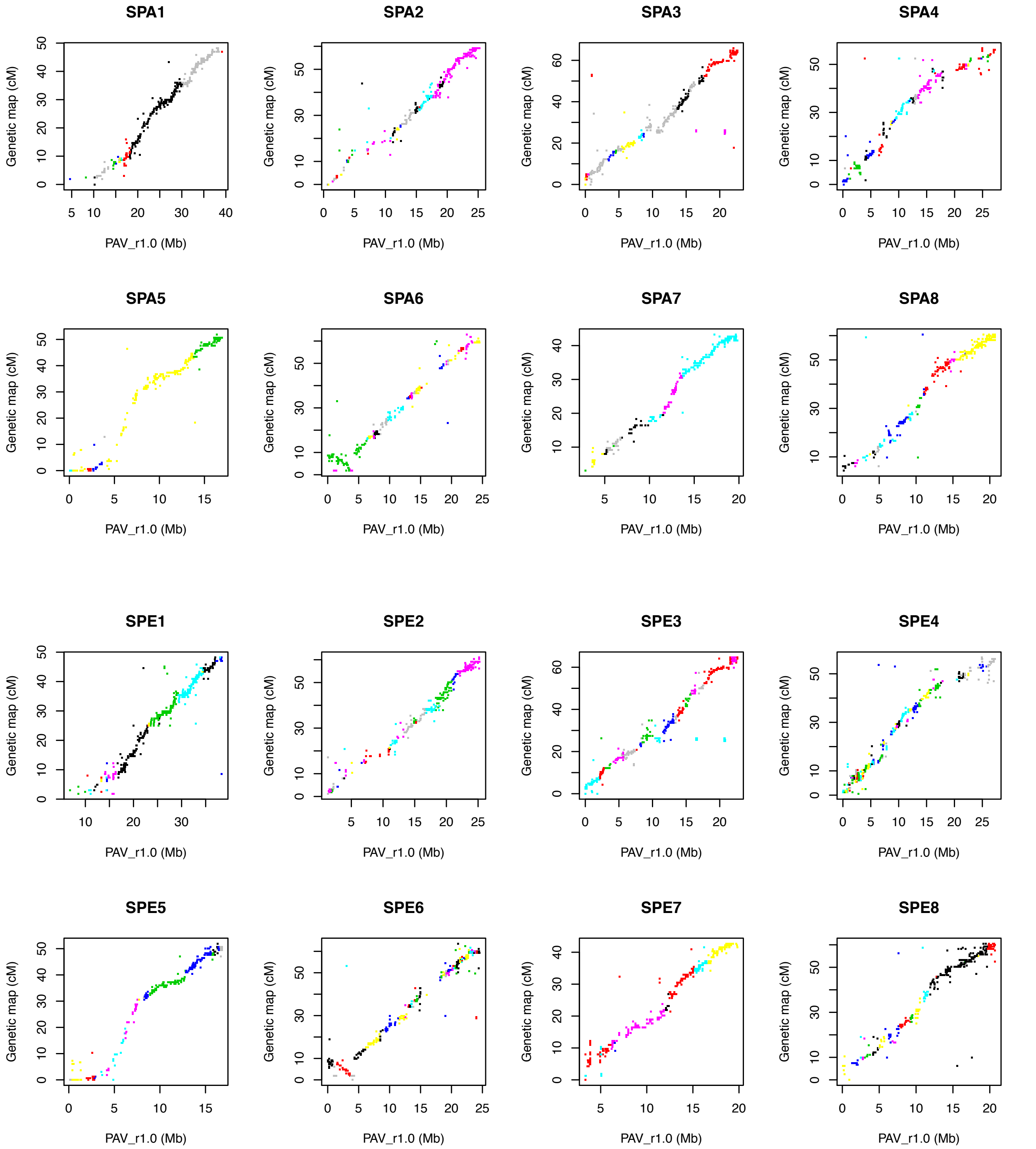


**Supplementary Figure S4.** Comparative analysis of the Somei-Yoshino genetic maps and the sweet cherry physical maps

Colors indicate contigs of the Somei-Yoshino genome sequences.

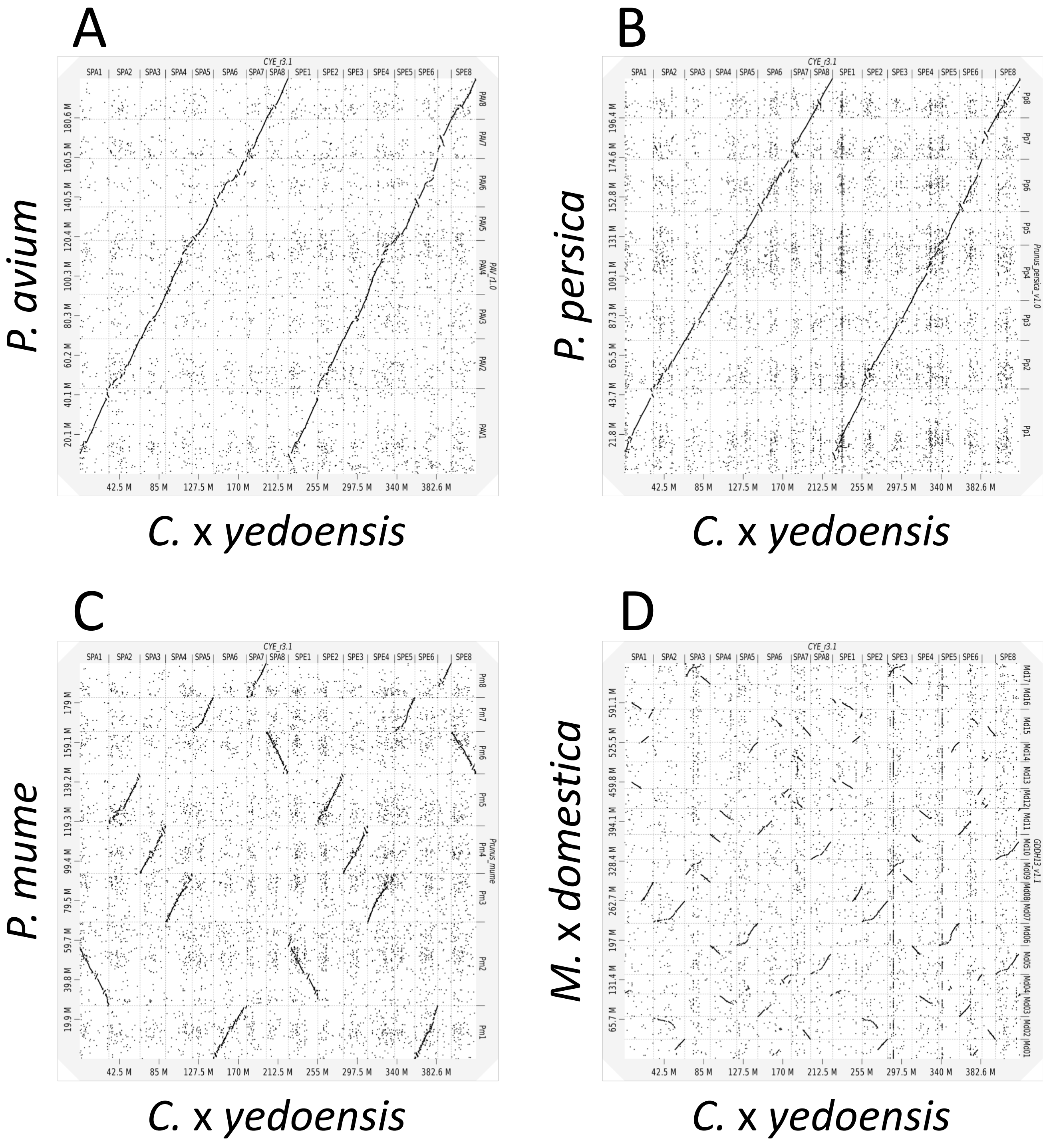


**Supplementary Figure S5.** Synteny of the Somei-Yoshino pseudomolecule sequences with those of taxa in the family Rosaceae

*X*- and *Y*-axes are sequences of Somei-Yoshino, *C.* × *yedoensis* (SPA1 to 8 and SPE1 to 8) and sweet cherry (A), peach (B), mume (C), or apple (D).

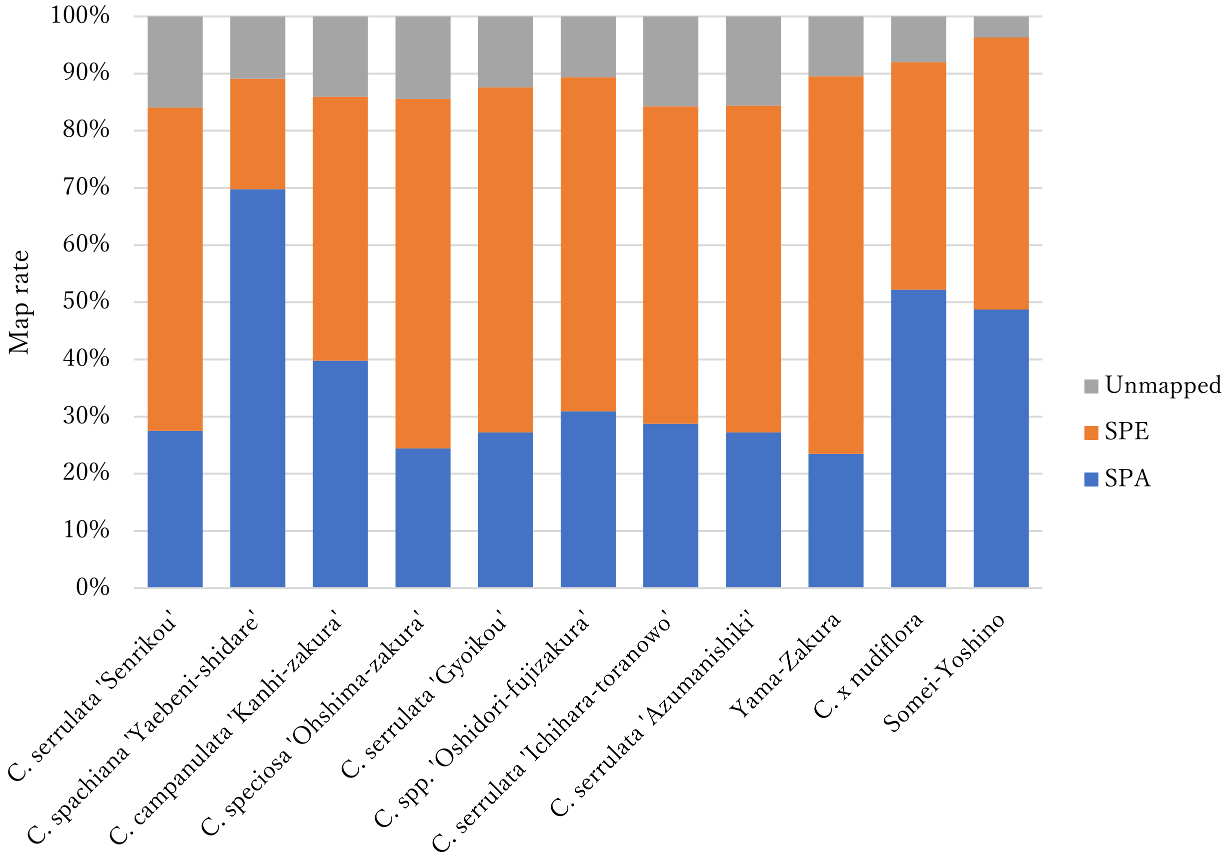


**Supplementary Figure S6.** Mapping rate of resequencing reads on the Somei-Yoshino genome sequence Reads are classified into three groups: mapped to CYE_r3.1spachiana (SPA) and CYE_r3.1speciosa (SPE) and unmapped.

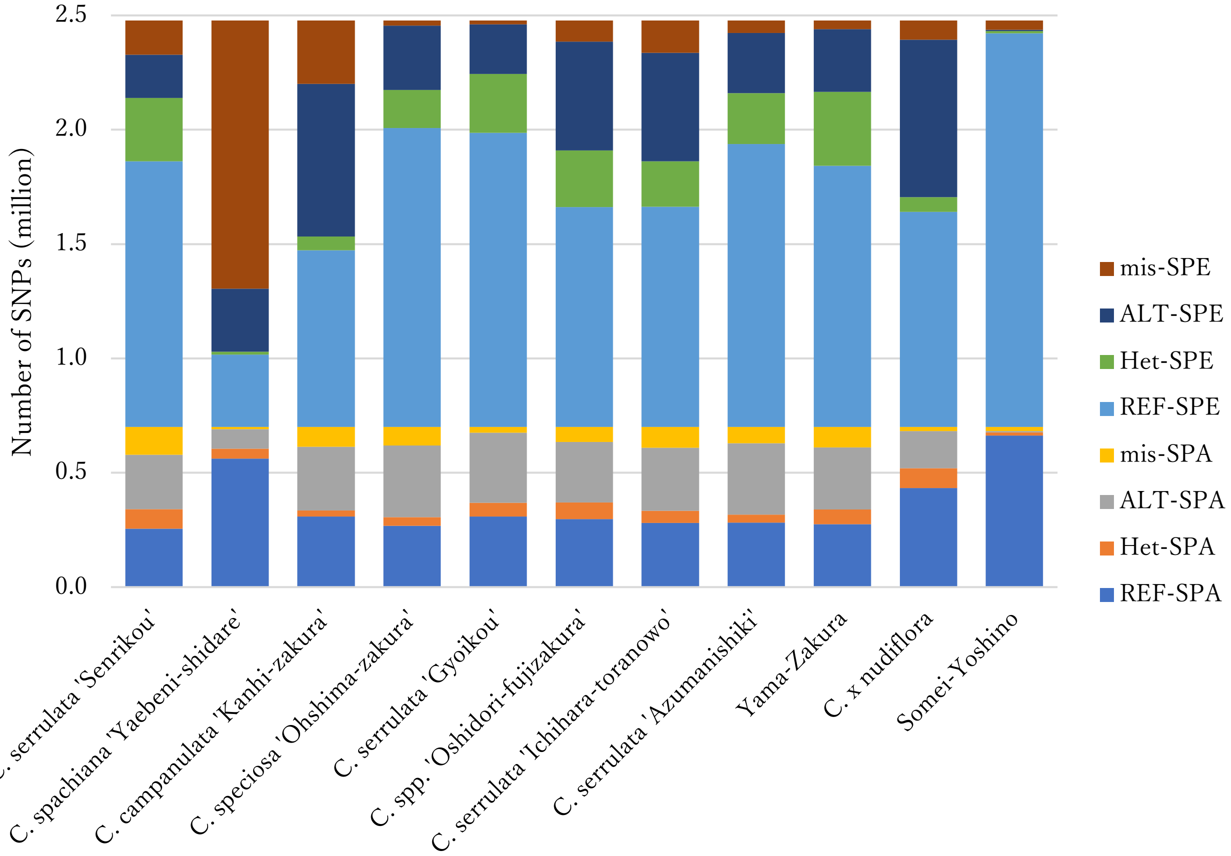


**Supplementary Figure S7.** Number of SNP genotypes with respect to Somei-Yoshino as a reference Homozygous SNPs as reference-type alleles and alternative-type alleles are indicated by REF and LAT, respectively. Heterozygous SNPs are denoted Het, and missing data are shown as mis. SNPs on CYE_r3.1spachiana and CYE_r3.1speciosa are indicated by SPA and SPE, respectively.

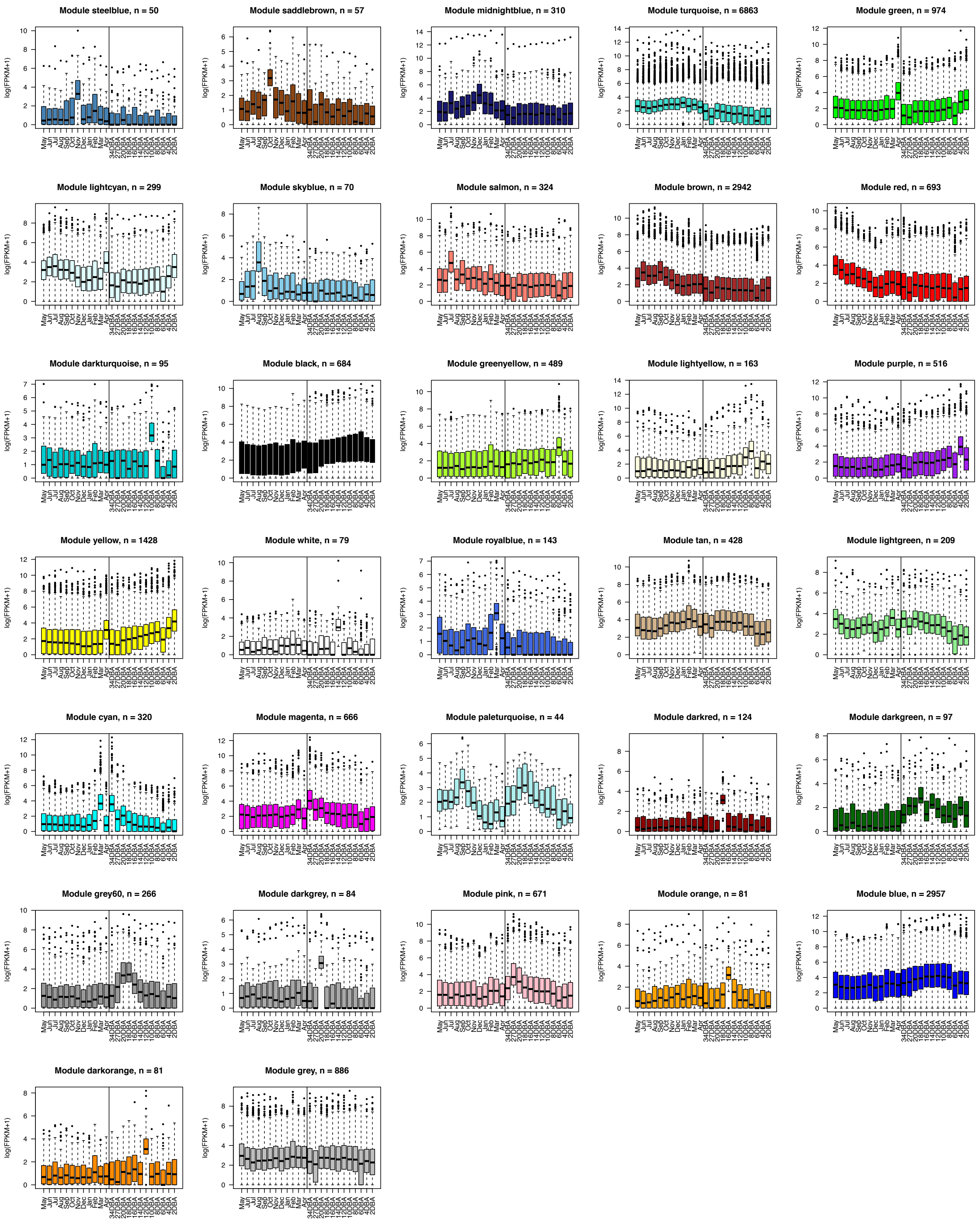


**Supplementary Figure S8.** Expression patterns of gene modules obtained from a weighted correlation network analysis

May to Apr are the months and 34DBA to 2DBA are days before anthesis when the bud samples were collected. Numbers of genes in each module are shown at the tops of boxplots.

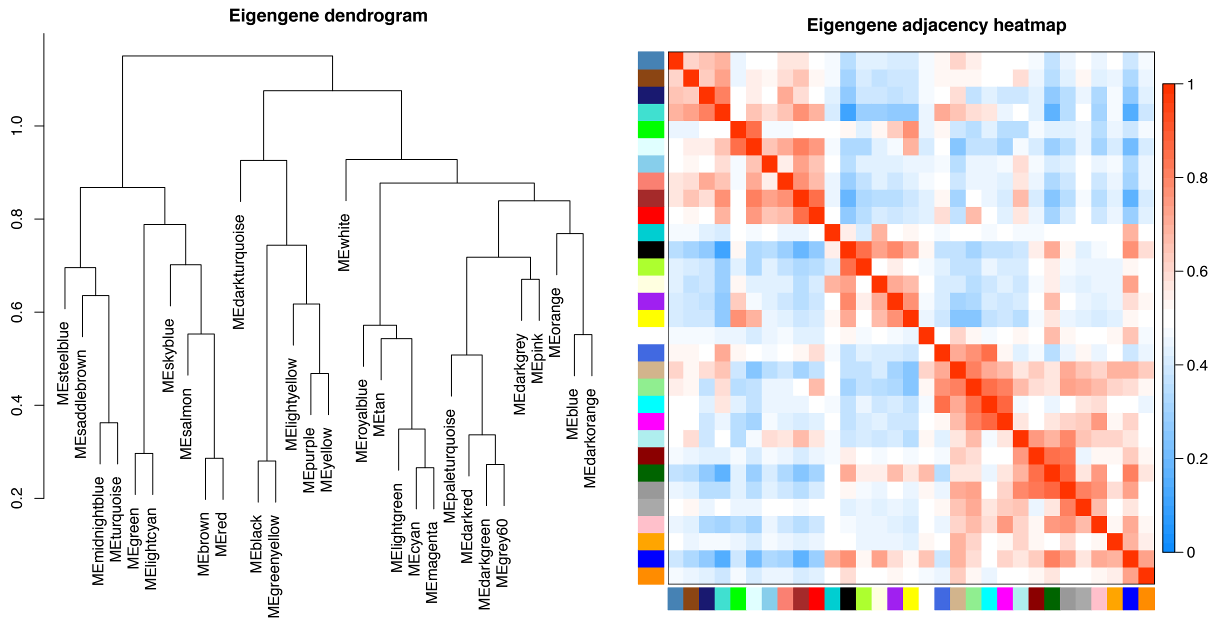


**Supplementary Figure S9.** Dendrogram and heatmap of gene modules represented by eigengenes obtained from a weighted correlation network analysis
